# Tissue-nonspecific alkaline phosphatase promotes neuronal cell proliferation and differentiation: metabolomic reveals glutathione and taurine as molecular correlates

**DOI:** 10.64898/2026.02.24.707745

**Authors:** Anne Briolay, Lionel G Nowak, Stéphane Balayssac, Veronique Gilard, David Magne, Caroline Fonta

**Affiliations:** Institute of Chemistry and Biochemistry (ICBMS), UMR 5246 CNRS-University of Lyon, Villeurbanne, France; University of Toulouse, CNRS, CerCo, Toulouse, France; University of Toulouse, CNRS, SOFMAT, Toulouse, France; Pathophysiology, diagnosis and treatments of musculoskeletal disorders lab (LYOS), UMR 1033 IN-SERM-Université de Lyon, Lyon, France

**Author notes:** Corresponding author: Lionel G Nowak, CerCo lab, UMR 5549, Pavillon Baudot, CHU Purpan, BP 25202, 31052 Toulouse Cedex, France.

## Abstract

Tissue-nonspecific alkaline phosphatase (TNAP) is a ubiquitous enzyme whose substrates are various phosphorylated extracellular molecules including pyridoxal phosphate (vitamin B6) and adenine nucleotides. Dysfunctions of TNAP result in hypophosphatasia, a rare disease characterized by defective bone mineralization and impaired brain functions. In the brain, TNAP expression peaks during development and is associated with various steps of neurogenesis. However, the influence of TNAP activity on neurogenesis remains poorly understood in its cellular and molecular aspects. Here we used the SK-N-SH D human neuroblastoma cell line as a cell culture model to further investigate the involvement of TNAP in neuronal precursor proliferation and neuronal differentiation. We also used ^1^H-NMR-based metabolomics to investigate the molecular correlates of TNAP action on SK-N-SH D cell proliferation and differentiation. We first observed an increase in alkaline phosphatase (AP) activity when the cells were placed in differentiation medium. We next found that inhibiting TNAP with a specific inhibitor (MLS-0038949) impeded neuroblastoma cell proliferation. TNAP inhibition also hindered neuronal differentiation, as evidenced by a decrease in the number of neurite-bearing cells. In contrast, neurite length was not affected by TNAP inhibition, suggesting that TNAP controls neurite sprouting, but not neurite outgrowth *per se*. The metabolomic results indicate that proliferation and differentiation are associated with a decrease in the amounts of proteinogenic amino acids as well as that of compounds potentially involved in lipid production. This analysis also revealed that proliferation and differentiation are associated with increased glutathione levels and decreased amounts of hypotaurine and taurine, supporting proposals that organosulfur compounds play an important role in these processes. Since pyridoxine was present in the culture media, these results suggest that TNAP is involved in neurogenesis through mechanisms in addition to its role in vitamin B6 metabolism and may instead involve the ectonucleotidase activity (or an unidentified activity) of TNAP.

## Introduction

Humans express four alkaline phosphatase (AP) isoenzymes (reviewed in Millán 2006). Two are expressed before birth in a tissue-specific manner (placental AP and germinal AP). One is expressed throughout life and is tissue-specific (intestinal AP). Diametrically opposed in its expression pattern, tissue-nonspecific alkaline phosphatase (TNAP, EC 3.1.3.1) is, as its name implies, expressed in several tissues such as bone, kidney, liver as well as in the brain both during development and in adulthood (Millán 2006; Fonta et al. 2015).

Historically mainly associated with bone formation and liver pathophysiology (Millán 2006), a large amount of data has been accumulated demonstrating that TNAP is also implicated in neuronal development as well as brain functions in adulthood. The first evidence for these roles derives from observations made in patients with dysfunctional TNAP, which results in a rare disease called “hypophosphatasia”. While the main clinical sign of hypophosphatasia is an alteration of skeletal mineralization (e.g., Rathbun 1948; Fraser 1957; Mornet et al. 2007; Whyte 2010), in its most severe forms (perinatal and infantile hypophosphatasia), it is also associated with pyridoxine-responsive epileptic seizures (e.g., Rathbun 1948; Sia et al. 1975; Baumgartner-Sigl et al. 2007; Taketani et al. 2014) and with abnormalities of the overall brain structure (Demirbilek et al. 2012; Hofmann et al. 2013; de Roo 2014; Fukazawa et al. 2018). In less severe forms, hypophosphatasia is accompanied by various neurological disorders such as fatigue and neuropathic pain (Colazo et al. 2019). TNAP knockout (KO) mouse models exhibit features of severe human hypophosphatasia, including epilepsy (Waymire et al. 1995; Narisawa et al. 1997). These mice further show reduced synaptogenesis and impaired myelination (Hanics et al. 2012).

Several lines of evidence indicate that TNAP is associated with brain development. For example, although significant TNAP activity is observed in the adult brain (e.g., Fonta et al. 2004; Langer et al. 2008; Negyessy et al. 2011), biochemical and histochemical studies have demonstrated that TNAP activity is at its highest during brain development (Goldstein and Harris 1981; Narisawa et al. 1994; Fonta et al. 2005; Brun-Heath et al. 2011). TNAP is also a marker of progenitor cells in the ventricular zone (VZ) and subventricular zone (SVZ) during embryonic development (Narisawa et al. 1994; Langer et al. 2007). In the adult brain, TNAP is also expressed in the SVZ of the lateral ventricles, one of the two neurogenic zones that persist in the adult (Langer et al. 2007; Kermer et al. 2010) (the neurogenic zone of the dentate gyrus in adult seems to be devoid of AP activity; Langer et al. 2007) and blockage of TNAP activity in adult SVZ reduces proliferation of stem cells and impairs the differentiation of progenitor cells into neurons and oligodendrocytes (Kermer et al. 2010). Yet the way TNAP controls the early stages of neuronal development remains little explored.

It is expected that the first step in this control relies on the regulation of extracellular levels of phosphorylated molecules. Yet TNAP is an ectoenzyme with little substrate specificity. Consequently, several extracellular phosphorylated compounds have been shown to be substrates of TNAP (Millán 2006; Whyte 2010; Zimmermann et al. 2012). In the brain, identified TNAP substrates are pyridoxal phosphate (PLP) (Waymire et al. 1995; Coburn 2015), adenine nucleotides (ATP, ADP, AMP) (Dorai & Bachhawat 1977; Ohkubo et al. 2000; Kaulich et al. 2003; Street et al. 2013) and phosphoethanolamine (Strickland et al. 1956). Other extracellular phosphorylated molecules (e.g., uridine nucleotides, phosphoproteins) could be hydrolyzed by TNAP, but formal evidence that this occurs *in situ* in the brain has not yet been provided. Cruz et al. (2017) identified several metabolites whose concentrations were significantly altered in the brain of TNAP KO mice; most of them (GABA, cystathionine, histidine for instance) are directly or indirectly linked to PLP-dependent enzymes while one (adenosine) may be linked to the ectonucleotidase activity of TNAP.

In this study, we used cultures of SK-N-SH D cells (Prehaud et al. 2020) to further explore the role of TNAP in the early stages of neuronal development. SK-N-SH D is a subclone of the SK-N-SH neuroblastoma cell line that can be induced to differentiate into catecholaminergic neurons (Biedler et al. 1973, 1978). Here, we took advantage of the presence of pyridoxine in the culture media to examine the function of TNAP during development, *apart from* its role in PLP metabolism. Using a specific TNAP inhibitor, we showed that TNAP participates in SK-N-SH cell proliferation and differentiation into neurons; on the other hand, although the appearance of neurites depended on TNAP activity, their growth *per se* did not appear to be affected by TNAP inhibition. To identify the metabolic pathways potentially involved in these actions of TNAP, we used an untargeted ^1^H-NMR-based metabolomics approach, which revealed significant variations in several important metabolites, most notably proteinogenic amino acids, taurine and glutathione, when proliferation and differentiation were hampered by TNAP inhibition.

## Materials and methods

### Cell culture and differentiation

Institutional ethical approval was not required for this study. The SK-N-SH D cell line (RRID: CVCL_B0GN; Pasteur Institute, Collection Nationale de Cultures de Micro-organismes I-5010; Prehaud et al. 2020) is a subclone of the SK-N-SH (RRID: CVCL_0531) human neuroblastoma cell line (Biedler et al. 1973). The SK-N-SH D cell line can differentiate at 100% as neuron-like cells in differentiating medium (Prehaud et al. 2020). The cell line was last authenticated by qPCR in 2005. The SK-N-SH D cell line is not listed as a commonly misidentified cell line by the International Cell Line Authentication Committee (ICLAC; http://iclac.org/databases/cross-contaminations/). The maximum number of passages was 22.

SK-N-SH D cells were grown in a proliferation medium at 37°C in a humidified atmosphere containing 5% CO_2_. The proliferation medium consisted in Dulbecco’s Modified Eagle Medium (Sigma-Aldrich, cat. no. D6546) containing 4.5 g/L glucose, 10% heat inactivated fetal bovine serum (FBS), 100 U/mL penicillin and 0.1 mg/mL streptomycin. In all experiments, cells were seeded in the proliferation medium at a density of 2,000 cells/cm^2^. For differentiation experiments, cells were first seeded in the proliferation medium and, after 24 h, the medium was replaced by a differentiation medium. The differentiation medium consisted in EndoGRO^TM^-LS complete medium from Millipore (cat. no. SCME001). This medium included ascorbic acid, rhEGF, hydrocortisone hemisuccinate, heparin sulfate and a reduced amount of FBS (2%) to induce differentiation into neurons. Day 0 corresponds to the day of replacement of the proliferation medium with the differentiation medium. Non-differentiated control cultures were grown in the proliferation medium for the same duration before harvesting.

### Protein concentration

Cell proteins were quantified using the BCA assay kit (Pierce, cat. no. 23227) after lysis of the cells by 0.2% Nonidet P40.

### Measurement of alkaline phosphatase activity

AP activity was examined after 1, 2 or 3 days in proliferation or differentiation media. Cells were scraped, washed with PBS and lysed in 0.2% Nonidet P40 (Euromedex, cat. no. UN3500-A). The samples were centrifuged and AP activity was determined in the supernatant using 10 mM p-nitrophenyl phosphate (pNPP, Sigma-Aldrich, cat. no. 71768) as substrate at pH 10.3 and at 37°C. AP activity was expressed relative to protein concentration. The experiment has been performed with 10 different batches of cells over 2 days (1 batch) or over 3 days (9 batches).

### TNAP inhibition

We used MLS-0038949 (TNAPi, Merck, cat. no. 613810) to inhibit TNAP activity. MLS-0038949 is a specific non-competitive inhibitor of TNAP (Sergienko et al. 2009; Dahl et al. 2009) with an *IC_50_* of about 0.2 µM (Sergienko et al. 2009; Dahl et al. 2009). This inhibitor has been used at 10-50 µM in electrophysiological studies of nervous tissue (Zhang et al. 2012; Street et al. 2013; Nowak et al. 2015; Choi et al. 2015; Gleizes et al. 2022) and cell culture studies (Ciancaglini et al. 2010; Huesa et al 2015; Bessueille et al. 2020).

We used MLS-0038949 rather than levamisole, the historic inhibitor of TNAP. Indeed, studies (Nowak et al. 2015) reported that levamisole, at the concentration traditionally used to inhibit TNAP (1-2 mM), also blocks action potential generation and therefore all activity-dependent processes. This makes levamisole unsuitable for live-cell studies. MLS-0038949 seems to be devoid of such non-specific actions as it does not alter action potential generation (Nowak et al. 2015) and postsynaptic response features (Gleizes et al. 2022).

We verified the efficacy of MLS-0038949 under our experimental conditions. For this purpose, we examined AP activity in the presence of 12 different concentrations (1 nM to 100 µM) of the inhibitor in two experiments. The dose-response relationship was fitted with Hill equation:

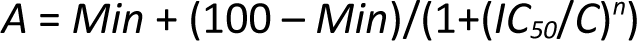

where *A* represent the normalized AP activity (in percent of AP activity in the absence of inhibitor), *Min* the maximal effect of the inhibitor (horizontal asymptote), “100” is the maximal AP activity (in the absence of inhibitor, 100%), *C* the inhibitor concentration, *IC_50_* the concentration producing half the maximal effect, and *n* the Hill slope. For the purpose of comparison, we performed in one experiment the same measurements using 8 concentrations (0.1 to 100 mM) of levamisole.

### Cell viability and metabolism in the presence of MLS-0038949

Cells were grown for 1 or 2 days in differentiation medium in the presence of 2 concentrations of MLS-0038949 (10 and 25 µM). The toxicity of MLS-0038949 was determined by measuring the activity of lactate dehydrogenase (LDH) released in the culture medium (Roche, Cytotoxicity Detection Kit, cat. no. 11644793001). LDH activity was expressed as a percentage of the value obtained in the presence of the TNAP inhibitor (MLS) minus that in control (Ctrl) normalized by the value obtained after lysis of the cells by Triton X100 (Trit) minus that in control (“% tox”): 100 × (MLS – Ctrl) / (Trit – Ctrl). A value of 0% corresponds to a toxicity that does not differ from that observed in control condition and a value of 100% corresponds to a toxicity level equivalent to that produced by the lysis of the cells with Triton X100. Each condition was realized in triplicate and the experiment was repeated four times.

Metabolic activity was determined by the tetrazolium salt method (MTT assay, Sigma-Aldrich, cat. no. M5655). Optical density measurements of formazan, produced by the reduction of MTT by mitochondrial dehydrogenases, were corrected by subtracting the optical density values obtained after cell lysis with Triton X100. Each condition was realized in triplicate and the experiment was repeated four times.

The same experiments have been performed on cells grown in the proliferation medium using 4 concentrations of MLS-0038949: 5, 10, 17.5 and 25 µM (supplementary Figure S1).

### Cell counting

After 1 or 2 days of differentiation in the presence or absence of MLS-0038949 (25 µM), cells were trypsinized, homogenized in phosphate buffered saline and counted on a Malassez counting chamber. The count was carried out in quadruplicate in four experiments.

### Assessment of differentiation and measurement of neurite length

Cells were grown in differentiation medium in the presence or absence of 25 µM MLS-0038949 . After 1 and 2 days, pictures were taken using an AxioVert 40 CFL microscope and an AxioCam ICc 1 camera (Zeiss). The experiment was repeated three times. The number of cell bodies and neurites were determined in 10 (in 2 experiments) or 20 (in 1 experiment) micrographs, resulting in 40 micrographs for each condition and a total of 160 scrutinized micrographs. Cells were considered differentiated when they produced at least one neurite whose length was at least twice the diameter of the cell body. The length of the neurites was measured with ImageJ software in the micrographs used to determine cell body and neurite numbers. Only cells considered differentiated were taken into account for these measurements. Occasionally two or three neurites issued from the same cell body could have a length > 2 cell body diameter and all were taken into account . When branching points were present, only the longest branch was measured.

### Proton NMR analysis

Five pairs of cultures have been used for metabolomics analyses. Each culture consisted in 5×10^6^ cells of the same batch of frozen cells. The cells were first seeded in proliferation medium for 24 h and then grown in differentiation medium for 2 days in the presence or absence of 25 µM MLS-0038949 . The cells were then washed on ice with cold phosphate buffered saline, scraped and then washed again twice before pelleting and freezing at -80°C.

The frozen cell pellets were extracted according to Beckonert’s procedure (Beckonert et al. 2007) using the methanol-chloroform-water system (2:2:1.425 (v/v/v)). At the first step of the procedure, duplicate protein assay was performed using the Bradford reagent (Bio-Rad, France) with bovine serum albumin as standard. Then, the upper aqueous phase was collected, dried and lyophilized. A deuterated borate buffer at pH=10.0 (60 mM), prepared in-house from borax (Merck, cat. no. S9640) and D₂O (Eurisotop, cat. no. D214B), was added to the dried aqueous extract with sodium 2,2,3,3-tetra-deutero-3-trimethylsilylpropionate (TSP, Merck, cat. no. 269913) as an internal chemical shift and quantification reference. The buffer also included 1 µM MLS-0038949 (in MeOH) to inhibit residual TNAP activity. The advantage of NMR chemical shift at pH 10 and the stability of metabolites under the chosen conditions has previously been demonstrated (Robert et al. 2011). ^1^H-NMR spectra were recorded at 298 K on a Bruker Avance 500 spectrometer equipped with a 5 mm TCI cryoprobe. A 1D pulse sequence (relaxation delay-pulse-acquisition) with pre-saturation for water (HOD) signal suppression was used. Acquisition parameters were as follows: relaxation delay 4.4 s, 30° pulse and acquisition time of 3.64 s. A spectral width of 18 ppm was used and 512 scans were collected. Data were processed using Bruker TopSpin software 3.2 with one level zero-filling and apodization (exponential, lb = 0.3 Hz) before Fourier transform, then phase correction, simple baseline correction and calibration (δ_TSP_ = 0 ppm) were applied. Metabolites were assigned with an in-house metabolite database at pH 10. For NMR data matrix, the bucket list was generated by the intelligent variable size bucketing tool included in the KnowItAll® package according to NMR signals assignment. Then the NMR - ProcFlow® pipeline (https://nmrprocflow.org/) was used for local baseline correction of spectra and alignment of NMR signals. Finally, the signals were integrated in accordance with the bucket list. Integrations were normalized by dividing the areas by that of the internal standard TSP and by the amount of proteins in the sample. In this work, the metabolomic approach was applied to assess relative metabolic variations across samples rather than for a comprehensive analysis using conventional multivariate statistical tools. A total of 36 metabolites were identified. Among them, adenosine was detected in a few samples but was excluded due to its very weak and noisy signal. The areas of the signals corresponding to the oxidized and reduced forms of glutathione were combined. A total of 34 variables were then considered for statistical analysis.

### Statistical analysis

The statistical analysis was made using Prism9/10 (GraphPad, Boston, MA, USA). Assessment of the normality of the data was conducted using Shapiro-Wilk test for n ≤ 5 or D’Agostino and Pearson test for n > 5. When the data were normally distributed, population data are presented as the mean and the 95% confidence interval of the mean is presented between brackets; otherwise, summary data are presented as the median and values between brackets indicate the 95% confidence interval of the median. The ROUT test was used to identify outliers, but only identified 3 out of ≈100 datasets. These outliers were not removed from the datasets. ANOVAs were used for AP activity, cell viability tests, protein content and cell proliferation. Following ANOVAs, paired comparisons were corrected for multiple comparisons by using Holm-Šídák’s multiple comparisons tests. Nested t-test was used for neurite length. For metabolomics results, the variable distributions did not depart from normality and all comparisons were made using two-tailed paired t-tests. A full statistical report is presented in the supplementary material (supplementary tables 1-8). The sample sizes were not predetermined statistically; however, the sample sizes used in the present study are comparable to those used in studies using similar approaches (e.g., Canals et al. 2005; Kermer et al. 2010; Díez-Zaera et al. 2011; Graser et al. 2015; Huesa et al 2015; Bessueille et al. 2020). Statistical power limitations are inherent to the small sample sizes, particularly for metabolomic data where n = 5 measurements per compound and per condition. However, the a posteriori power appears reasonably high for most metabolites whose quantities varied following TNAP inhibition (Supplementary Table 8).

## Results

### Alkaline phosphatase activity increases in differentiating SK-N-SH D cells

Using pNPP as substrate, we first examined whether SK-N-SH D cells displayed AP activity and whether this activity depended on time in culture and culture media. Cells were cultured for 1 to 3 days. AP was present across all days and conditions. However, we found that AP activity level depended on both the culture medium and the number of days in culture (Figure 1A). The number of days in culture had an overall mild effect on AP activity (two-way ANOVA, *P* = 0.05, Figure 1A). Post hoc tests (Holm-Šídák’s multiple comparisons tests) showed that the effect of days in culture was significant only in the differentiation medium with a significantly higher AP activity on day 3 compared to day 1 (*P* = 0.02, Figure 1A).

**Figure 1.**
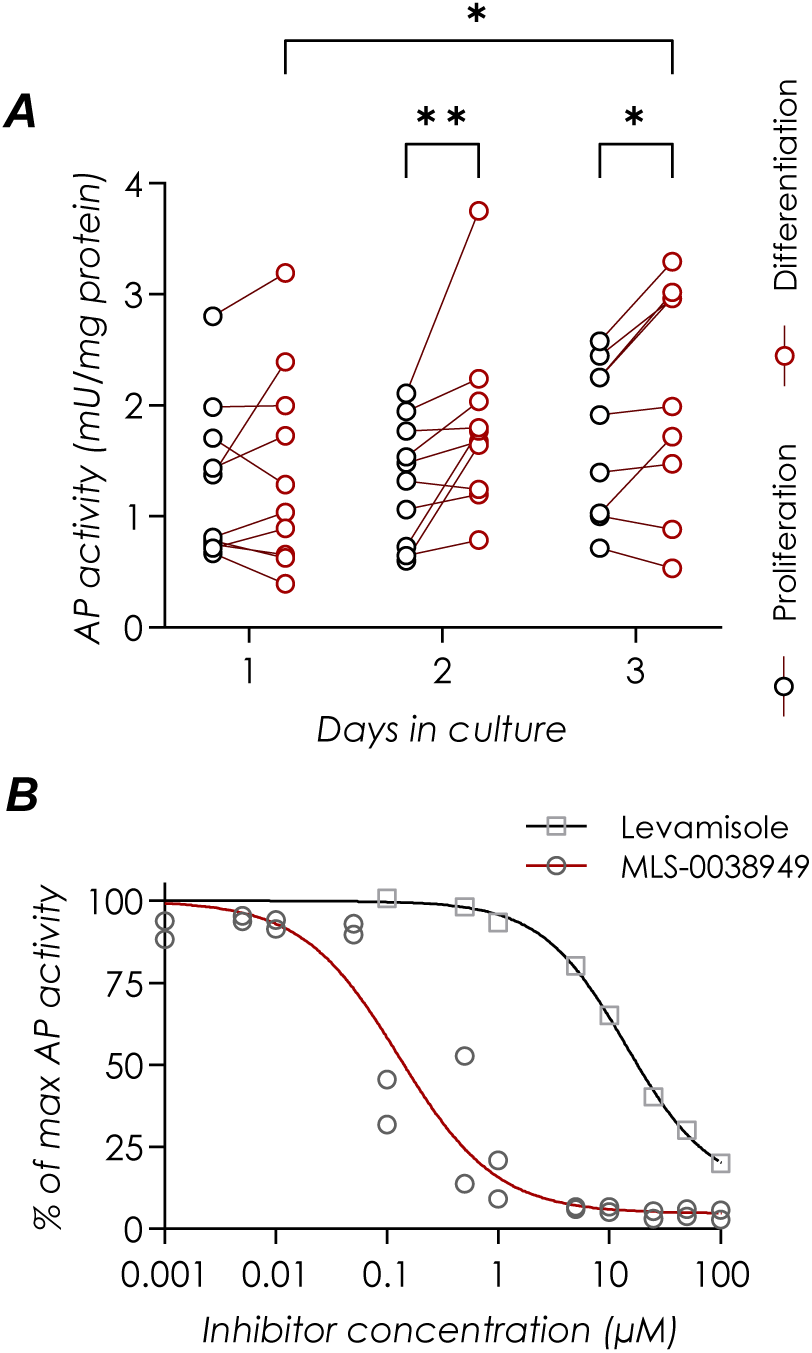
Alkaline phosphatase activity increased when SK-N-SH D cell cultures were grown in differentiation medium and was suppressed in the presence of TNAP inhibitors. Alkaline phosphatase (AP) activity was determined using p-nitrophenyl phosphate (pNPP, 10 mM) as substrate. **A.** Effect of culture medium. The cells were grown for 1, 2 or 3 days either in proliferation medium or in differentiation medium (day 1 and day 2: n = 10 cultures for each medium; day 3: n = 9 cultures for each medium). Each dot corresponds to one culture. Dots joined by lines correspond to pairs of cultures that were seeded at the same time in either proliferation or differentiation medium (black and red dots respective-ly). AP activity was normalized by the protein amount. Two-way ANOVA (mixed-effect analysis) revealed a significant effect of culture media on AP activity (*P* = 0.004) and a weaker effect of time in culture (*P* = 0.05). Post hoc tests (Holm-Šídák’s multiple comparisons test) showed that the effect of culture medium was significant after 2 or 3 days in culture and that AP activity was higher after 3 days in culture in proliferation medium than after 1 day in culture in proliferation medium. Significance levels of post hoc tests are given by the number of asterisks: *: *P* < 0.05; **: *P* < 0.01. B. Inhibition of TNAP activity by MLS-0038949 (n = 2 experiments) and levamisole (n = 1 experiment) after two days in differentiation medium. Twelve concentrations of MLS-0038949 (1 nM to 100 µM) and eight concentrations of leva-misole (100 nM to 100 µM) have been tested. AP activity (dots) was normalized by the activity observed in the absence of inhibition (max activity, 100%). The dose-response relationships were fitted with Hill equation (continuous lines). The *IC_50_* returned by the fit were 0.13 µM (95% CI: 0.08-0.20 µM) for MLS-0038949 and 17 µM (95% CI: 16-19 µM) for levamisole. The Hill slope (*n*) was close to -1 (MLS-0038949: -0,9930; levamisole: -1,006). The remnant activity observed with the highest concentrations of MLS-0038949 (25-100 µM) was at 4.7% of the activity in the absence of inhibitor (horizontal asymptote).

Culture medium had a stronger effect on AP activity. AP activity was higher in the differentiation medium compared to proliferation medium (two-way ANOVA, *P* = 0.004). The effect of culture medium was not significant on day 1 (Holm-Šídák’s multiple comparisons test, *P* = 0.4) but there was a significant increase in AP activity in differentiation medium compared to proliferation medium at both day 2 (*P* = 0.002) and day 3 (*P* = 0.03) (Figure 1A). The medians of AP activity ratios in differentiation vs. proliferation media were 118.4% [101.8% - 243.6%] on day 2 and 121.2% [88.5% - 133.9%] on day 3 . In summary, we observed that AP activity was approximately 20% higher in the differentiation medium compared to the proliferation medium.

To determine whether TNAP was responsible for this AP activity, we examined the action of two specific TNAP inhibitors: levamisole (van Belle 1972) and MLS-0038949 (Sergienko et al. 2009; Dahl et al. 2009). Dose-response relationships (Figure 1B) were established in two experiments for MLS-0038949 and in one experiment for levamisole. AP activity was determined as above from cells grown for 2 days in differentiation medium. Both inhibitors nearly completely suppressed all AP activity, although with different efficacy: the *IC_50_* obtained from Hill equations fitted to the data was much lower with MLS-0038949: 0.13 µM, vs. 14.2 µM for levamisole (Figure 1B). These *IC_50_* values are in close agreement with previously reported values (e.g., MLS-0038949: Sergienko et al. 2009; Dahl et al. 2009; levamisole: van Belle 1976; Debray et al. 2013). The residual activity observed with the highest concentrations of MLS-0038949 (25-100 µM) corresponds to an asymptote at 4.7% of the activity without inhibitor. This implies that MLS-0038949 was able to suppress >95% of the AP activity exhibited by SK-N-SH D cells.

### Inhibition of tissue-nonspecific alkaline phosphatase activity reduces cell proliferation

The increase in AP activity in differentiation medium led us to examine whether TNAP is involved in cell proliferation, in neuronal differentiation and in neurite growth. For this purpose, we inhibited TNAP activity using MLS-0038949 while cells were grown in differentiation medium.

To examine the effect of inhibiting TNAP on cell proliferation, cells were grown for one or two days in the presence or absence of MLS-0038949 at 25 µM. The number of cells has been determined in quadruplicate in four cultures for each condition and each culture duration. As illustrated in Figure 2A, there was an effect of both time (two-way ANOVA, *P* < 0.0001) and TNAP inhibitor (*P* = 0.02) on the number of cells per well. First, there was an increase in the number of cells with time. This increase was significant for both control cultures (+147%; *P* < 0.0001, Holm-Šídák’s multiple comparisons test) and for cultures grown in the presence of MLS-0038949 (+64%; *P* = 0.004).

**Figure 2.**
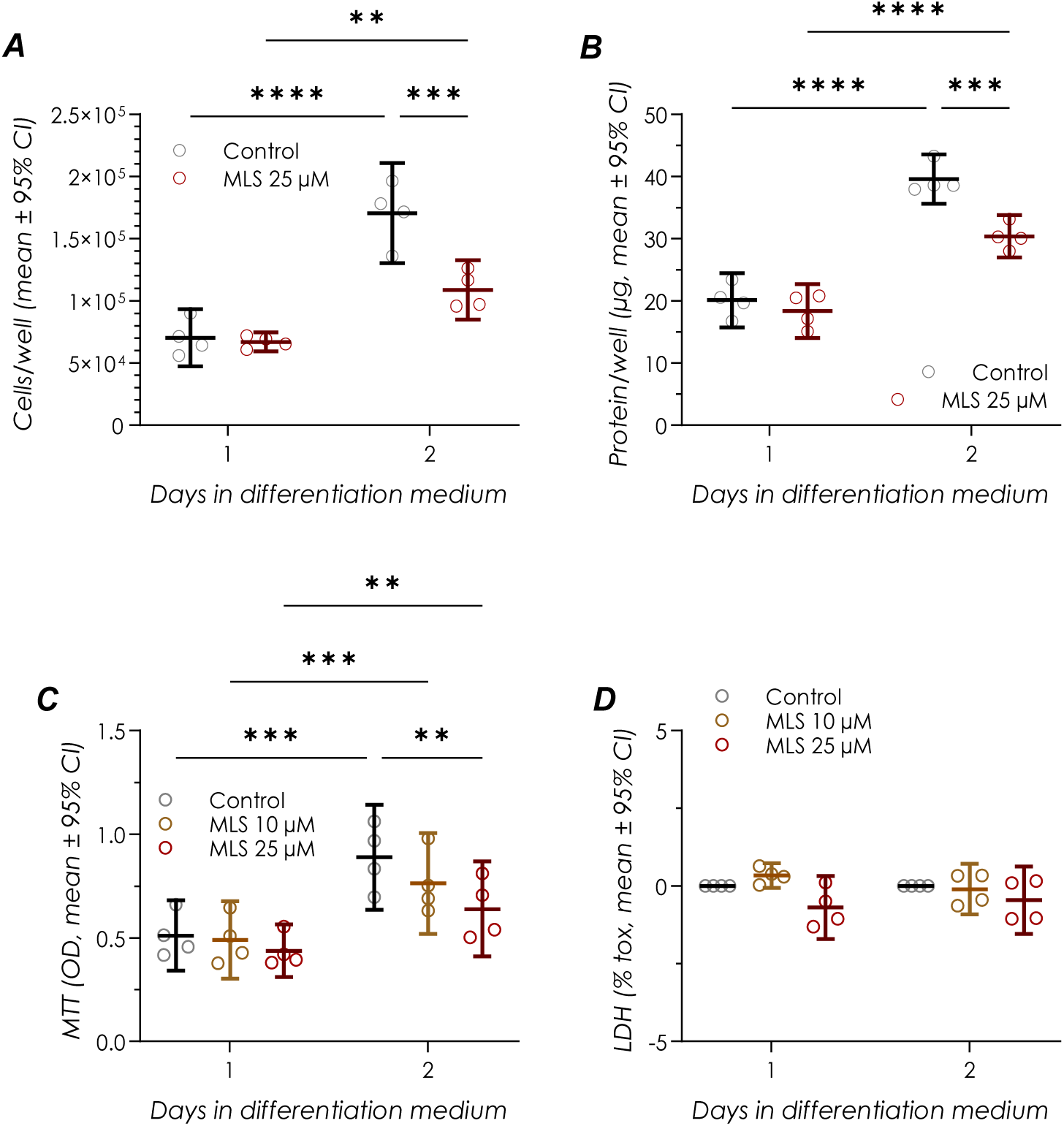
Effects of inhibiting TNAP activity with MLS-0038949 on cell number, protein amount, metabolic activity and cell viability. Cells were grown for one or two days in the differentiation medium in the presence or absence of the TNAP inhibitor MLS-0038949 (MLS). Dots in the graphs correspond to single experiments. The central horizontal line corresponds to the means and the upper and lower bars encompass the 95% confidence interval of the mean. All comparisons were made using two-way ANOVA with time and MLS treatment as fac-tors and post hoc comparisons of pairs of means were all performed using Holm-Šídák’s multiple comparisons test. Significance levels of post hoc tests are given by the number of asterisks: **: *P* < 0.01; ***: *P* < 0.001; ****: *P* < 0.0001. **A.** Cell proliferation was reduced by the TNAP inhibitor (n = 4 pairs of cultures for each day). The number of cells per well increased with time both in control cultures and in cultures grown in the presence of MLS-0038949 (25 µM). However, inhibition of TNAP activity with MLS-0038949 resulted in a significant de-crease of cell proliferation visible in cultures grown for two days. **B.** Changes in protein amount (protein per well) with time and with MLS-0038949 (25 µM) treatment parallel those observed for cell numb er (n = 4 pairs of cultures for each day). **C.** Cell metabolic activity was assessed using the MTT assay and two concentrations of MLS-0038949 (10 and 25 µM). There was an overall effect of time on formazan level. MLS-0038949 had no significant effect in cultures grown for 1 day. MLS-0038949 at 25 µM significantly reduced the overall formazan level in cultures grown for 2 days, reflecting the decrease in cell number observed in the same condition (n = 4 triplets of cultures for each day). OD: optical density of formazan. **D.** The possibility that MLS-0038949 exerts a cytotoxic effect was determined by using the LDH release assay (n = 4 triplets of cultures for each day). Two MLS-0038949 concentrations were used (10 and 25 µM). LDH activity values were normalized such that a value of 100% corresponds to the toxicity induced by the complete cell lysis produced by Triton X100, and 0% to the toxicity measured in control condition. Geisser-Greenhouse correction was used to correct for lack of sphericity for the two-way ANOVA. The results show an overall but weak protective effect of MLS-0038949 (ANOVA, *P* = 0.048) but this does not show up in pairwise comparisons.

Second, this analysis revealed that inhibiting TNAP reduced cell proliferation. This effect of MLS-0038949 was not significant after one day in culture (*P* = 0.8) but was significant on day 2 (*P* = 0.0004). On day 2, MLS-0038949 treatment resulted in a reduction in cell number to 66% of the control value (Figure 2A).

The amount of protein was determined in the same experiments as for cell counts . As illustrated in Figure 2B, changes in protein amount closely paralleled changes in cell number. There was an effect (two-way ANOVA) of both time in culture (*P* < 0.0001) and MLS-0038949 (25 µM) treatment (*P* = 0.02). Unsurprisingly given the increase in cell number, the amount of protein increased with time both in control cultures (+99% between day 1 and day 2; *P* < 0.0001, Holm-Šídák’s multiple comparisons test) and in cultures grown in the presence of MLS-0038949 (day 2 vs. day 1: +67%; *P* < 0.0001). Besides, TNAP inhibition led to a reduction in protein amount compared to control on day 2 (*P* = 0.0005), although not on day 1 (*P* = 0.4). On day 2, the protein amount was reduced to 77% of that of the control cultures. As a consequence of the parallel changes in cell number and protein content, the amount of protein per cell was quite similar between control and treated cultures after one day in proliferation medium (means: 0.29 and 0.27 ng/cell respectively) and after two days in proliferation medium (means: 0.24 and 0.28 ng/cell respectively).

Cell metabolism was evaluated using the MTT test and two concentrations of MLS-0038949 (10 and 25 µM). Cultures were grown in the presence or absence of the inhibitor for 1 or 2 days in the same four experiments as above. As shown in Figure 2C, there was again an effect of both time and MLS-0038949 on formazan absorbance level (two-way ANOVA, *P* = 0.002 for time and *P* = 0.003 for treatment). Post hoc tests (Holm-Šídák’s tests) indicated an increase in formazan level as a function of time (control: *P* = 0.001; MLS-0038949 10 µM: *P* = 0.0004; MLS-0038949 25 µM: *P* = 0.001). The increases in formazan levels from day 1 to day 2 was +61% in control cultures and +46% in cultures treated with MLS-0038949 at 25 µM. These changes were in the same direction, although less marked, as the increases in cell number from day 1 to day 2 (Figure 2A). After one day in culture, MLS-0038949 (25 µM) had no influence on formazan level (*P* = 0.2) but it reduced it to 72% of control after two days in culture (*P* = 0.002, Figure 2C). This value is quite similar to the reduction in cell number observed on day 2 in the presence of MLS-0038949 (Figure 2A). Thus, as with protein amount, the reduction in overall metabolic activity likely reflects the decrease in cell proliferation resulting from TNAP inhibition.

The results presented above point toward parallel effects of TNAP inhibition on cell number, protein amount, and metabolism. To ascertain that these changes were not due to a toxic effect of MLS-0038949, we examined cell viability using the LDH release assay in the same cultures as for MTT tests. Time in culture had no significant effects on LDH release (two-way ANOVA, effect of days in culture: *P* = 0.7) (Figure 2D). Opposite to that expected from a toxic effect, MLS-0038949 application resulted in a small reduction in LDH release that was weakly significant (two-way ANOVA, effect of treatment: *P* = 0.05). However, changes in LDH release were not significant anymore when tested in pairwise fashion (Holm-Šídák’s tests, *P* ≥ 0.2). Overall, these results indicate that the decrease in cell number induced by inhibiting TNAP (Figure 2A) was not due to an increase in cell death and was rather the consequence of a reduction in the rate of cell proliferation.

MTT assays and LDH activity measurements have also been performed in preliminary experiments on cells in proliferation medium, with a similar outcome (supplementary Figure S1): MLS-0038949 also reduced overall metabolic activity but had no toxic effect.

### Inhibition of tissue-nonspecific alkaline phosphatase activity impedes neuronal differentiation

In addition to its dramatic effect on cell proliferation, we next found that inhibiting TNAP also strongly affected neuronal differentiation. As above, SK-N-SH D cells were cultured in differentiation medium for 1 or 2 days. Three pairs (with or without MLS-0038949 25 µM) of cultures were examined. In differentiation condition, cells could be subdivided in two categories: those bearing neurites, *i.e.*, neurons, and those that did not grow appreciable neurites, *i.e.*, non-differentiated cells or weakly differentia-ted cells (considered as such when neurite lengths were less than twice the cell body diameter) (Figure 3A-C). Most differentiated cells showed only one neurite longer than the criteria of twice the cell body diameter, but occasionally two or three neurites could be seen emerging from the same cell body. The number of neurites per cell was calculated for 40 micrographs for each day and condition . In each picture, the number of neurites was counted and divided by the number of cell bodies; the greater the number of differentiated cells, the closer the ratio is to 1 (Figure 3D). The number of days in culture had a significant effect (two-way ANOVA, effect of days in culture: *P* = 0.02) on the number of neurites per cell, but it vanished in post hoc comparisons (Holm-Šídák’s tests, day 1 vs day 2: control: *P* = 0.1; MLS-0038949: *P* = 0.2).

**Figure 3.**
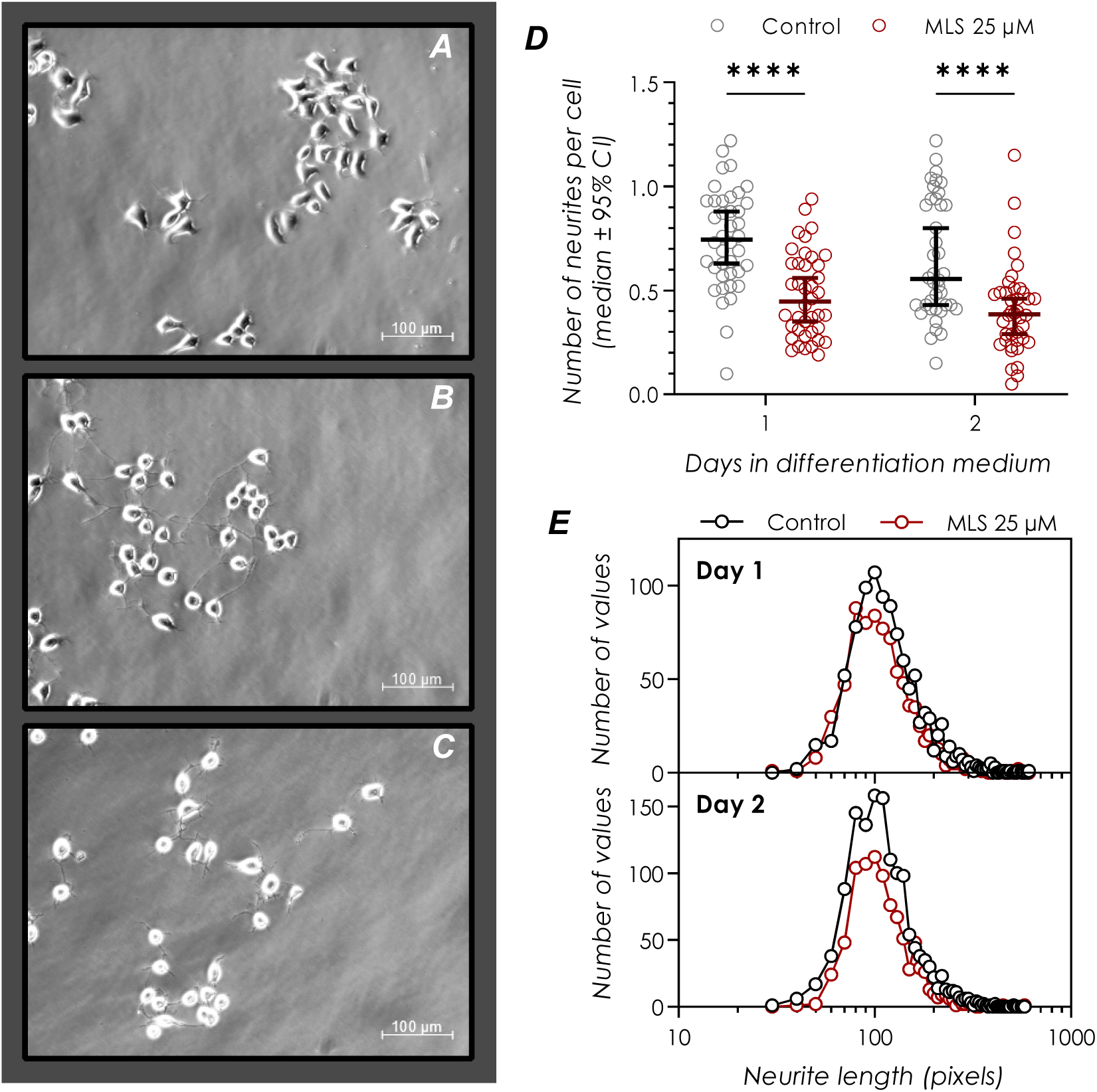
Inhibition of TNAP reduces the proportion of differentiated cells but does not affect the length of the neurites of differentiated cells. **A-C**: Micrograph of SK-N-SH D cells maintained in: proliferation media for one day (A), differentiation media for two days (B) and differentiation media with MLS-0038949 (25 µM) for two days (C). **D.** TNAP is involved in the differentiation of SK-N-SH D cells into neurons. Cells were considered differentiated when they bore at least one neurite whose length was twice the cell body diameter. The number of neurites was then normalized by the number of cells in each micrograph . Each dot in the scattergram corresponds to the number of neurites per cell determined for one micrograph. Forty micrographs have been analyzed for each day and condition. The micrographs taken on day 2 were not taken on the same cells and neurites as those of day 1. The central horizontal line corresponds to the median of the number of neurites per cell and the upper and lower bars encompass the 95% confidence interval of the median. The total number of cells that have been classified as either differentiated or not differentiated was: day 1, control: 1398; day 1, MLS-0038949 25 µM: 1689; day 2, control: 2157; day 2, MLS-0038949 25 µM: 2257. Two-way ANOVA and Holm-Šídák’s post hoc tests revealed a strong effect of the TNAP inhibitor. ****: *P* < 0.0001. **E.** Distribution of neurite length of differentiated cells measured after one or two days in differentiation medium in the presence or absence of MLS-0038949 (25 µM). The length of the neurites of differentiated cells was not affected by TNAP inhibition. Overall, neurite length has been measured for 4124 differentiated cells in 4 × 40 micrographs (day 1, control: 1017 neurons; day 1, MLS-0038949: 824 neurons; day 2, control: 1379 neurons; day 2, MLS-0038949: 904 neurons).

On the other hand, MLS-0038949 treatment resulted in a marked reduction in the number of neurites per cell (ANOVA and Holm-Šídák’s post hoc tests: *P* < 0.0001, Figure 3D). On day 1, cells bore 0.75 (median) neurite [0.63 - 0.88, *n* = 40 pictures] in control condition but only 0.45 [0.35 - 0.56, *n* = 40 pictures] in the presence of MLS-0038949 – a decrease of 40%. On day 2, the median of the number of neurites per cell was 0.56 [0.43 - 0.80, *n* = 40] in control condition but was only 0.39 [0.29 - 0.46, *n* = 40] when TNAP was inhibited – a decrease of 30%. These results suggest that TNAP contributes to neuronal differentiation of SK-N-SH D cells.

### Neurite length is not affected by inhibition of tissue-nonspecific alkaline phosphatase in differentiated neurons

While inhibition of TNAP reduced the proportion of cells bearing neurites, it did not affect the length of the neurites of these differentiated cells. Neurite lengths have been measured in the same cultures and micrographs as for the proportion of differentiated cells presented above. Distributions of neurite length were unimodal. As summarized in Figure 3E, there was no significant effect of TNAP inhibition on neurite length (nested t-test on log transform of the data; day 1: *P* = 0.3; day 2: *P* = 0.9). Medians of neurite length were: day 1, control: 120 pixels [116-124]; day 1, MLS-0038949: 114 pixels [100-118]; day 2, control: 112 pixels [109-114]; day 2, MLS-0038949: 110 pixels [107-114]. Thus, although TNAP promoted the appearance of neurites, it did not appear to intervene significantly in their growth once it had been initiated.

### Metabolome

The results presented above positively identified a role of TNAP in cell proliferation and in neuronal differentiation. To identify molecules reflecting and potentially mediating these actions of TNAP, we used a metabolomics approach. Metabolomics is the comprehensive study of small molecules, or metabolites, providing a global snapshot of cellular metabolic processes. In this study, metabolomic analyses were performed using ^1^H-NMR, which enables a non-targeted metabolic profiling without prior knowledge of the detected metabolites. The metabolome was analyzed in five pairs of cultures, with one member of the pair receiving no treatment (control) and the other receiving MLS-0038949 (25 µM). The cells were harvested after two days in differentiation medium. In these cultures too, MLS-0038949 treatment resulted in a marked decrease (paired t test, *P* = 0.006) in protein amount to 76% [65% - 86%] of the control value (not illustrated). The relative abundances of metabolites have thus been normalized by the amounts of proteins, which reflects the number of cells as mentioned above. Figure 4A presents a typical ^1^H-NMR spectrum of the aqueous extract from a control SK-N-SH D cell culture. Thirty-six metabolites have been detected by ^1^H-NMR. Of these, we excluded adenosine due to its weak and noisy signal while we combined the signals corresponding to oxidized and reduced glutathione. The signals of the metabolites whose amounts varied significantly with MLS-0038949 treatment (see below) are presented at higher magnification in Figure 4A (in boxes or indicated by ar-rows).

**Figure 4.**
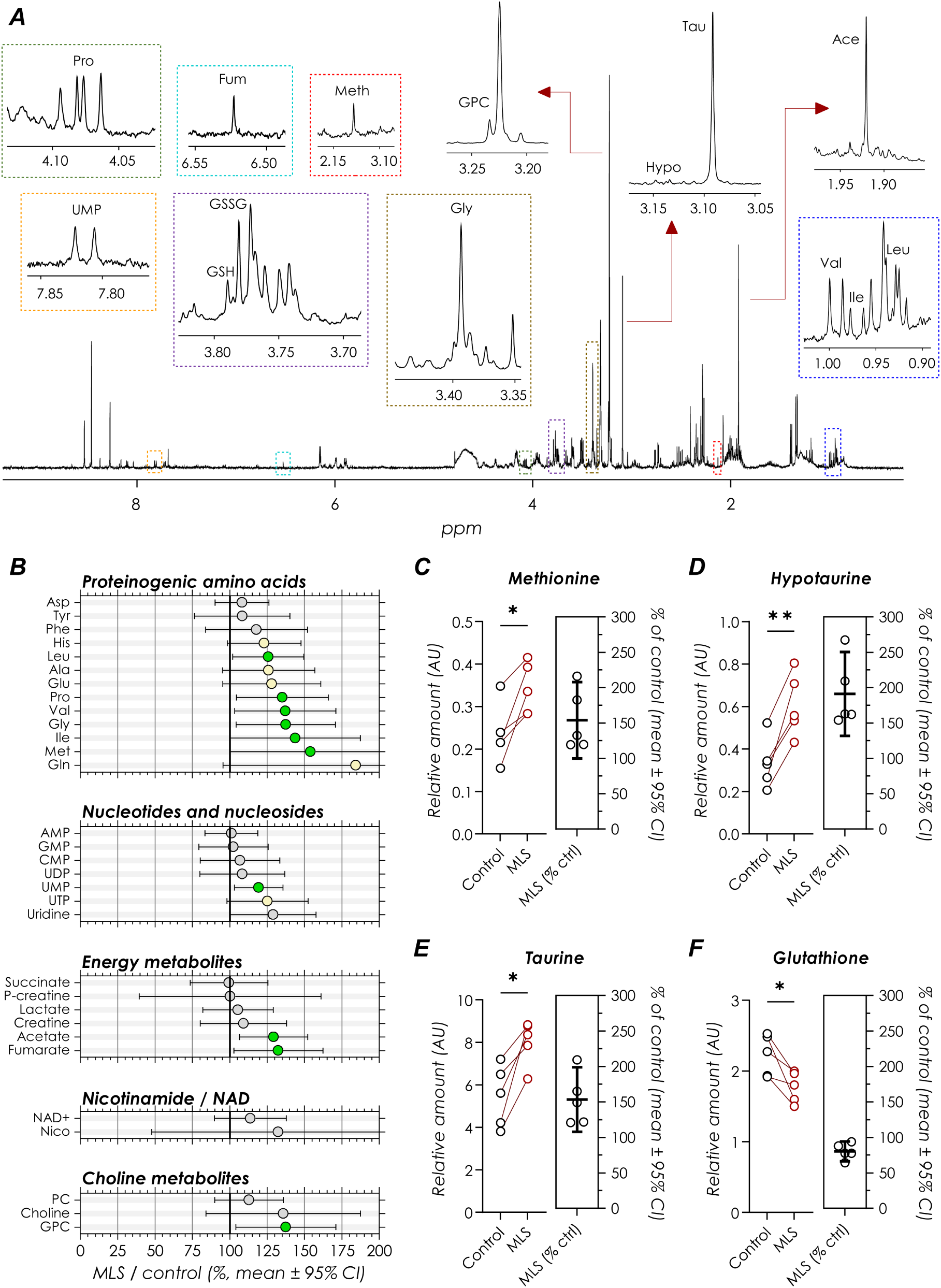
Metabolome of SK-N-SH D cells after two days in differentiation medium in the presence or absence of the TNAP inhibitor MLS-0038949 (25 µM). **A.** ^1^H-NMR spectrum of the aqueous extract from an SK-N-SH D untreated cell culture after two days in differentiation medium. The zoomed-in sections, highlighted in colored dashed boxes or pointed at by arrows, correspond to the signals of the metabolites showing the most significant variations (green dots in B and metabolites in C-F): proline (Pro, green box), UMP (orange box), fumarate (Fum, cyan box), methionine (Met, red box), oxidized and reduced glutathione (GSH and GSSG, purple box), glycerophosphocholine (GPC), glycine (Gly, brown box), hypotaurine and taurine (Hyp and Tau), acetate (Ace), and valine, isoleucine and leucine (Val, Ile and Leu, blue box) . **B.** Ratios of the concentrations expressed as percentages for 31 of the 34 metabolites identified in aqueous extracts of SK-N-SH D cells cultured for two days in differentiation medium in the presence of MLS-0038949 *vs*. control (not shown: hypotaurine, taurine and glutathione which are detailed in panels D-E) (n = 5 pairs of cultures) . Symbols correspond to the mean ratio and bars encompass the 95% confidence interval of the mean. Green symbols correspond to metabolites whose amounts changed significantly in the presence of MLS-0038949 (paired t-test, *P* < 0.05), yellow symbols to marginally significant differences (0.05 ≤ *P* ≤ 0.06) and gray symbols to no significant changes. PC: phosphocholine. **C-F.** Focus on changes in amounts for organosulfur compounds. Each dot in scattergrams on the left corresponds to one culture. Dots joined by lines correspond to pairs of cultures that were seeded at the same time in proliferation medium with (red dots) and without (black dots) MLS-0038949 (n = 5 pairs of cultures). Right scattergrams: ratio of concentration in MLS vs. control (mean and 95% confidence interval indicated by central and flanking lines). The level of methionine (C), hypotaurine (D) and taurine (E) increased significantly while that of glutathione (F) decreased in the presence of MLS-0038949. Significance levels of paired t-tests are given by the number of asterisks: *: *P* < 0.05; **: *P* < 0.01. The probability of making a type 1 error being set at *P* = 0.05, either one or two of the significant differences reported here would be a false positive (0.05 × 34 = 1.7). Yet significant differences were observed in a much larger number of comparisons (13/34 comparisons).

Thirteen of the 20 proteinogenic amino acids were detected (Figure 4B). The level of 6 of these increased significantly (*P* < 0.05, paired t-test, green dots in Figure 4B) in cultures treated with MLS-0038949: methionine (mean increase: +54% [100% - 208%]), isoleucine (+44% [100% - 187%]), glycine (+37% [104% - 171%]), valine (+37% [103% - 171%]), proline (+35% [104% - 166%]) and leucine (+26% [102-150%]). Concentration for four amino acids showed an increase that stood on the margin of the significance level (0.05 ≤ *P* ≤ 0.06, yellow dots in Figure 4B): glutamine (+84% [95% - 273%]), glutamate (+28% [95% - 161%]), alanine (+26% [95% - 157%]) and histidine (+23% [98% - 148%]). Three amino acids (tyrosine, phenylalanine and aspartate) showed no significant variation (gray dots in Figure 4B). There were no amino acids whose concentration would decrease significantly.

Seven nucleotides/nucleosides have been identified (Figure 4B): AMP, CMP, GMP, uridine, UMP, UDP and UTP. Of these, two uridine nucleotides increased or tended to increase in the presence of MLS-0038949: UMP (*P* = 0.02, +19% [103% - 135%]) and UTP (*P* = 0.051, +25% [98% - 152%]).

Six metabolites related to energy metabolism were retrieved (Figure 4B). Creatine and phosphocreatine levels were not affected by TNAP inhibition. Lactate and succinate levels also showed no variation. Only acetate and fumarate amounts increased in cultures treated with the TNAP inhibitor (acetate: *P* = 0.01, +29% [106% - 152%]; fumarate: *P* = 0.04, +32% [103% - 162%]).

MLS-0038949 treatment had no effect on the levels of nicotinamide and nicotinamide adenine dinucleotide (NAD^+^) (Figure 4B).

^1^H-NMR allowed detecting choline and two related metabolites (Figure 4B). Inhibiting TNAP had no significant effect on the level of choline and phosphocholine, but the amounts of glycerophosphocholine increased by more than 35% in cultures treated with MLS-0038949 relative to control ( *P* = 0.02, +37.5% [104% - 171%]).

The amounts of all the organosulfur compounds detected by ^1^H-NMR showed significant variation with MLS-0038949 treatment. These results are detailed in Figure 4C-F. Methionine level increased (*P* = 0.02, +54% [100% - 208%], Figure 4C), as well as that of hypotaurine (*P* = 0.003, +91% [132% - 250%], Figure 4D) and taurine (*P* = 0.01, +53% [109% - 199%], Figure 4E). Glutathione, on the other hand, demonstrated a significant decrease (*P* = 0.03, 80% of control [66% - 94%], Figure 4F). Thus, cells that differentiated more (control condition) contained more glutathione and less taurine and hypotaurine than cells whose differentiation was hindered by TNAP inhibition.

## Discussion

Our study demonstrated the presence of AP activity in both proliferating and differentiating SK-N-SH D cells. This AP activity was nearly completely suppressed by MLS-0038949, a specific TNAP inhibitor. We also observed that AP activity increased when cells were bathed in differentiation medium. Inhibition of TNAP during the differentiation process had two main effects on neurogenesis: it reduced cell proliferation and it partially blocked neuronal differentiation, as revealed by a reduction in the proportion of cells bearing neurites. Yet, for the subpopulations of cells that grew neurites, neurite length was not affected by TNAP inhibition. These results suggest that the proliferation of precursor cells and their transitions to neurons are controlled by extracellular compounds that acquire their efficacy only after dephosphorylation by TNAP. In parallel, metabolomic results showed that inhibiting TNAP during differentiation resulted in an increase in concentration for most amino acids and strongly affected organosulfur compounds level. These results suggest that differentiation, under the control of TNAP, is associated with a change in the fate of sulfur-containing amino acids towards glutathione rather than taurine.

### TNAP and alkaline phosphatase activity in neurogenesis

To the best of our knowledge, our study is the first that detected AP activity in the SK-N-SH neuroblastoma cell line before differentiation. Previous studies using the SH-SY5Y cell line, a subclone of the SK-N-SH cell line, either reported (Kohring and Zimmermann 1998) or did not observe (Graser et al. 2015) significant AP activity before differentiation. Our result is also in line with numerous *in situ* histochemical studies that showed AP activity on neurons’ precursor cells, especially at the level of proliferative zones in embryos (ventricular and subventricular zones) and at the level of the neurogenic zone of the lateral ventricle in the adult (Narisawa et al. 1994; Mishra et al. 2006; Langer et al. 2007; Kermer et al. 2010; Brun-Heath et al. 2011).

We observed that AP activity increased when cells were placed in differentiation medium. Using SH-SY5Y cells, Graser et al. (2015, their supplementary data) observed an increase in expression of the mRNA for TNAP once the cells were placed in differentiation medium, although the difference from t0 became significant only on the 8^th^ day. Díaz-Hernández et al. (2010) reported presence of TNAP activity in differentiated cells of the same cell line. Using the P19 teratocarcinoma cell line, Scheibe et al. (1991) showed that retinoic acid, which is classically used to induce neuronal differentiation, also induced an increase in TNAP mRNA expression and a 3 fold increase in AP activity, which was maximal 3 days after induction of differentiation. Using the 1C11 cell line, Ermonval et al. (2009) and Brun-Heath et al. (2011) showed that TNAP expression and AP activity appeared during neuronal differentiation, whereas these were not detected in undifferentiated cells. Finally, using primary cultures of hippocampal neurons, Díez-Zaera et al. (2011) showed an increase in TNAP protein expression and AP activity as a function of time that paralleled neuritogenesis.

Our study goes beyond the mere observation of a correlation between the presence of TNAP and proliferation or differentiation. Indeed, we showed that TNAP plays a prominent role both in the proliferation and the differentiation of SK-N-SH D cells. First, proliferation appeared to be controlled by TNAP as inhibiting TNAP reduced the number of cells after two days in culture by 32% in comparison to control. Using a different model – cultured neuronal stem cells derived from the SVZ of the adult mouse –, Kermer et al. (2010) also showed that proliferation was strongly reduced when TNAP was inhibited with shRNA. Apart from the brain, a recent study showed that TNAP deficiency also impairs proliferation of bone and muscle progenitor cells (Zhang et al. 2021).

Second, differentiation into neurons, witnessed by the sprouting of neurites, was impeded by inhibiting TNAP. In control condition, most cells grew neurites once in the differentiation medium. Yet inhibition of TNAP resulted in a lower proportion (−30% to −40%) of differentiated cells both after 1 and 2 days in culture. Thus, neuronal differentiation is partially determined by TNAP activity. In cell cultures derived from adult SVZ progenitors, Kermer et al. (2010) reported an impairment of differentiation (evidenced by reduction in expression of doublecortin) when TNAP expression was blocked by shRNA . Besides, our results appear to be complementary to those of Graser et al. (2015), who found that overexpression of TNAP in SH-SY5Y cells increased the expression of neuronal markers such as NEUN, TAU or NPY. Apart from the brain, blockage of TNAP activity has also been reported to compromise adipogenesis and differentiation of osteoblast precursor cells (reviewed in Estève et al. 2016).

Yet, once neurite growth was initiated, we observed no difference in neurite length between control and treated cultures. This may appear to be at variance with the study by Díez-Zaera et al. (2011), who observed that inhibiting TNAP reduced *axonal* growth in primary cultures of cortical or hippocampal neurons. This discrepancy may be due to the use of different models (mouse primary neuron vs human neuroblastoma culture). This could also result from the use of different TNAP inhibitors, as some of the observations by Díez-Zaera et al. (2011) were based on the use of levamisole. However, it is worth noting that Díez-Zaera et al. (2011) also reported that TNAP inhibition had no effect on the growth of the *dendrites*. Therefore, there would be no discrepancy between our study and theirs if the neurites examined in the present study — after 1 or 2 days in differentiation medium — were dendrites rather than axons. This possibility is supported by studies performed on differentiating SH-SY5Y cells. In these cells, the expression and immunolabeling of microtubule-associated protein 2 (MAP2), which is specific to dendrites, are present from the first days of differentiation (Gimenez-Cassina et al. 2006; Graser et al. 2015; Paik et al. 2019); on the other hand, axon markers such as tubulin associated unit protein (tau protein) or medium molecular weight neurofilament protein (NF 160) are only expressed and visible in neurites after several days (generally ≥ 5) of differentiation (Encinas et al. 2000; Gimenez-Cassina et al. 2006; Graser et al. 2015; Paik et al. 2019).

### Extracellular phosphorylated compounds possibly regulated by TNAP

Our results implicate that proliferation and differentiation of SK-N-SH D cells are regulated by molecules whose extracellular levels depend on TNAP. It is well established that TNAP plays a crucial role in the metabolism of PLP (reviewed in Coburn 2015), a form of vitamin B6 that acts as a cofactor for numerous enzymes (e.g., Percudani and Peracchi 2009). PLP cannot cross cell membranes and its de-phosphorylation by TNAP to pyridoxal is a prerequisite for its intracellular incorporation (reviewed in Spector and Johanson 2007; Coburn 2015). Consistent with this function of TNAP, analysis of the metabolome of TNAP KO mice showed concentration changes for several compounds whose synthesis or degradation depend directly or indirectly on PLP-dependent enzymes (Waymire et al. 1995; Cruz et al. 2017).

Yet our results are unlikely to be due to faulty vitamin B6 metabolism, because the culture media were supplemented with (non phosphorylated) pyridoxine. Amount of pyridoxine (pyridoxine HCl) in the proliferation solution was 4 mg/L, which corresponds to a concentration of 19.5 µM: a value several tens of times higher than the concentration of vitamin B6 in the cerebrospinal fluid ( ≈0.1-0.4 µM: Spector 1978; Albersen et al. 2014). The amount of pyridoxine in the differentiation solution is not disclosed by the supplier, but we expect it to be in the same range as in the differentiation medium. Thus, perturbation of PLP dependent enzymatic processes would not explain our results. In support to this contention, the increase in taurine and decrease in glutathione concentration we observed are opposite to those observed in vitamin B6 deficiency: vitamin B6 deficiency in rats led to an *increase* in glutathione level in the liver (Lima et al. 2006) while severe vitamin B6 restriction in rats resulted in a huge *decrease* (to less than 1% of the control level) of urinary taurine (Sturman and Cohen 1971); a decrease in taurine amount has also been reported in the brain of 14 day old rats deprived of vitamin B6 (Guilarte 1989). This does not mean that the metabolism of PLP by TNAP is unimportant for normal brain development (e.g., Cruz et al. 2017), but it does mean that in addition to PLP, other extracellular compounds regulated by TNAP also play a role in this process.

Another possible extracellular substrate for TNAP corresponds to membrane phosphoproteins. Although Fedde et al. (1993), using fibroblasts from hypophosphatasic patients, concluded that “ALP does not modulate the phosphorylation of plasma membrane proteins”, studies using other preparations isolated a couple of proteins that could constitute TNAP substrates: Sarrouilhe et al. (1992) found that liver TNAP is able to dephosphorylate an unidentified 18 kDa phosphoprotein; using P19 teratocarcinoma, Scheibe et al. (2000) demonstrated that TNAP is able to dephosphorylate an unidentified 98 kDa protein; Ermonval et al. (2009) reported that laminin (∼400 to ∼900 kDa) is maintained in a weakly phosphorylated state by TNAP in the cell line 1C11. Overall, there seem to be only few phosphoproteins that could be dephosphorylated by TNAP *in situ*, and their identity and roles in neurogenesis remain largely unknown.

Apart from PLP and phosphoproteins, other extracellular substrates for TNAP are adenine nucleotides. Action of TNAP as an ectonucleotidase has been reported many times in various organs and tissues (reviewed in Millán 2006; Zimmermann et al. 2012) as well as in the brain (Dorai & Bachhawat 1977; Street et al. 2013) and in the neuroblastoma × glioma cell line NG108-15 (Ohkubo et al. 2000; Kaulich et al. 2003). This ectonucleotidase activity is also supported by the decrease in adenosine level in the brain of TNAP KO mice (Cruz et al. 2017). Thus the changes we observed during TNAP inhibition may result from either an increase in extracellular ATP concentration or from a decrease in concentration of the extracellular ATP degradation down-products, in particular adenosine . While we did not attempt to quantify the extracellular levels of ATP and adenosine, it is worth mentioning these because of the well-established roles of purinergic signaling in different steps of neurogenesis (reviewed in: Zimmerman 2011; Grković et al. 2019; Rimbert et al. 2023). Interestingly, using SH-SY5Y cultures, which is the model closest to ours, Canals et al. (2005) demonstrated that adenosine acting through both A1 and A2_A_ receptors is involved in inducing neuronal differentiation. Using different cell lines, Abbracchio et al. (1989) (IMR32 neuroblastoma cell line) and Charles et al. (2003) (PC12 cell line) also reported that A2_A_ adenosine receptor activation promoted neuronal differentiation.

Studies have shown that neurite growth proper also depends on purinergic signaling; however, depending on the models and structures studied, different purine receptors have been involved, some promoting and others inhibiting neurite growth (Thevananther et al. 2001; Cheung et al. 2005; Canals et al. 2005; Díez-Zaera et al. 2011; Rubini et al. 2014; Heine et al. 2015; Ribeiro et al. 2016; Alçada-Morais et al. 2021; Mut-Arbona et al. 2023; Díez-Zaera et al. 2024). In our experimental conditions, neurite length was neither increased nor decreased in the presence of the TNAP inhibitor. This suggests either that purinergic signaling was not involved during the first two days of neurite growth of SK-N-SH D cells, or, as several ectonucleotidases are involved in brain development (reviewed in Goding et al. 2003; Zimmermann and Langer 2015; Grković et al. 2019), that the control of neurite growth proper depended on ectonucleotidases other than TNAP.

### Metabolome

#### Amino acids and proteins

The overall protein content was reduced by inhibition of TNAP (Figure 2B). This lower protein content most likely resulted from the decrease in the number of cells since the protein content normalized by the number of cells appeared to be fairly constant across conditions. MTT tests also showed that TNAP inhibition led to a decrease in global metabolic activity, in a proportion similar to the reduction in cell number and protein content.

Metabolome analysis allowed identifying 13 proteinogenic amino acids (Figure 4B). Almost all showed a significant (*P* < 0.05) or marginally significant (0.05 ≤ *P* ≤ 0.06) increase in concentration when TNAP was inhibited. Increases in concentration ranged from +23% (histidine) to +84% (glutamine). Since the protein content per cell appears to be fairly constant across all conditions, this accumulation of amino acids suggests that protein turnover was reduced in the presence of the TNAP inhibitor.

#### Phospholipid and lipid production

Inhibiting TNAP resulted in a reduction of the proportion of cells bearing neurites. It can be expected that this reduction was accompanied by an overall reduction of cell membrane area, hence a reduced need for lipid incorporation. This reduced need for lipids is possibly corroborated by the increase in the levels of three metabolites related to lipid metabolism: acetate, uridine nucleotides and glycero-phosphocholine.

We noticed an increase in the amount of acetate in the cells bathed with the TNAP inhibitor. Acetate can be condensed with coenzyme A to generate acetyl CoA; acetyl CoA, in turn, is essential for fatty acid synthesis (Wakil et al. 1983). Although acetate is involved in many metabolic pathways, its accumulation in cells whose differentiation was slowed by TNAP inhibition may reflect reduced lipogenesis in this condition.

Uridine nucleotides are linked, though indirectly, to glycerophospholipid synthesis. Glycerophospholipids constitute one of the two main classes of lipids in brain cell membranes (the other being cholesterol). The major subclasses of glycerophospholipids are phosphatidylethanolamines and phosphatidylcholines, which represent ≈ 20% and > 30% of all brain lipids (Wells and Dittmer 1967; Fitzner et al. 2020). Interestingly, studies showed that phosphatidylcholine synthesis is strongly stimulated in SK-N-SH cells induced to differentiate (Cook et al. 1996). This results in a 40% increase in phosphatidylcholine amount *per cell* (Marcucci et al. 2010). Similar results were also reported with the PC12 cell line (Araki and Wurtman 1997). The major synthesis pathway for phosphatidylcholines and phosphatidylethanolamines (Kennedy’s pathway) uses CTP, either in combination with choline to generate CDP-choline for the synthesis of phosphatidylcholines, or in combination with ethanolamine to generate CDP-ethanolamine for the synthesis of phosphatidylethanolamines (van der Veen et al. 2017). CTP, which is maintained at low concentration within cells, was not detected in the present study. Yet CTP is generated from UTP (Richardson et al. 2003; Pooler et al. 2005) and studies showed that uridine nucleotides metabolism is intimately linked with that of cytidine nucleotides and with CDP-choline production (Richardson et al. 2003; Cansev 2006). Of the 7 nucleotides/nucleosides that we have detected, two showed significant or nearly significant increases in amounts in the presence of the TNAP inhibitor: UMP (*P* = 0.02) and UTP (*P* = 0.051). This suggests that uridine nucleotide consumption, and indirectly that of CTP, was reduced under these conditions. This would be consistent with reduced glycerophospholipid synthesis when differentiation was hindered by TNAP inhibition.

Finally, we observed an increase (+37.5%) in the amount of glycerophosphocholine during TNAP inhibition. Glycerophosphocholine is a down product of phosphatidylcholine degradation. The increase in the amount of glycerophosphocholine that we reported would correspond to an increased turnover of phosphatidylcholine consecutive to reduced incorporation of phosphatidylcholines into cell membranes when differentiation was impeded by TNAP inhibition.

Altogether, the increase in the amounts observed for acetate, glycerophosphocholine and uridine nucleotides in the presence of the TNAP inhibitor suggests reduction in lipogenesis that could reflect the reduction in the number of neurites per cell.

#### Glutathione vs. hypotaurine and taurine

Glutathione level decreased (−20%) significantly when TNAP activity was inhibited. Among the identified metabolites, glutathione was actually the only one whose concentration decreased in the presence of MLS-0038949. Reduction of glutathione level goes against a possible oxidative stress induced by MLS-0038949. Long term effect (≥ 24 h) of oxidative stress (if it does not kill all the cells right away) is a large increase in glutathione level (Shimizu et al. 2002; Tirmenstein et al. 2005), as well as an upregulation of the enzymes involved in glutathione synthesis (Tirmenstein et al. 2005). Besides, the results of our LDH assays indicate that MLS-0038949 was devoid of toxic effects.

Glutathione is a tripeptide composed of glutamate, cysteine and glycine. The availability of these 3 glutathione precursors is unlikely to explain the reduction of glutathione level we observed when TNAP was inhibited: glycine level was elevated (+37%, *P* < 0.05), as well as that of glutamate (+28%, 0.05 ≤ *P* ≤ 0.06) and glutamate precursor glutamine (+84%, 0.05 ≤ *P* ≤ 0.06) in the presence of MLS-0038949. Cysteine was not detected in our ^1^H-NMR measurements, presumably because it is maintained at low concentration to prevent its endogenous cytotoxicity (Janáky et al. 2000; Stipanuk et al. 2006). However, its concentration also probably increased as the amount of methionine, cysteine principal precursor (Peck and Awapara 1967; Kwon and Stipanuk 2001), was significantly increased (+54%, *P* < 0.05); besides, a shortage of cysteine would not allow the large increase in hypotaurine and taurine concentration we observed (see below).

Instead, the *increase* in the amount of sulfur amino acids induced by TNAP inhibition may explain the decrease in glutathione level. Glutathione is synthesized by two enzymes (reviewed in Dringen and Hirrlinger 2003; Stipanuk et al. 2006; Lu 2013): glutamate-cysteine ligase (GCL, EC 6.3.2.2) and glutathione synthetase (EC 6.3.2.3). GCL is the rate-limiting enzyme. GCL activity is regulated by both cysteine and methionine availability, but counter-intuitively: when their concentrations *increase*, GCL activity *decreases* (Ohta et al. 2000; Kwon and Stipanuk 2001). It is when levels of cysteine and methionine are low that GCL expression is induced and that its activity is increased (Stipanuk et al. 2006). Therefore, the increase in methionine concentration we observed, and that of cysteine concentration that likely results from it, should reduce GCL activity and glutathione production.

Reversing perspectives, our results suggest that glutathione level increased when cells proliferated and differentiated without constraint (i.e., in control conditions). Glutathione is well known for its roles as an antioxidant and in detoxification. Furthermore, studies have shown that cell proliferation is associated with increased glutathione levels and that glutathione is in fact causally involved in cell cycle progression (reviewed in Pallardó et al. 2009; Lu 2013). In addition to proliferation, neuronal differentiation may also be associated with increases in glutathione level. In particular, using the SH-SY5Y cell line, Takahashi et al. (2016) observed that differentiation promoting insulin or IGF-1 induced an increase in glutathione amounts. There is therefore reason to hypothesize that TNAP inhibition led, through molecular mechanisms and signaling pathways that remain to be established, to a decrease in glutathione synthesis, this decrease then slowing down the proliferation and/or differentiation of SK-N-SH D cells.

In contrast to glutathione, we found that inhibition of TNAP induced a strong increase in hypotaurine (+91%) and taurine (+53%) amounts. Taurine is synthesized from cysteine in 3 enzymatic steps (reviewed in Tappaz et al. 1992; Stipanuk et al. 2006; Ramírez-Guerrero et al. 2022): cysteine dioxygenase (CDO, EC 1.13.11.20) converts cysteine to cysteine sulfinic acid, which is decarboxylated to hypotaurine by cysteine sulfinate decarboxylase (EC 4.1.1.29); hypotaurine is converted by hypotaurine dehydrogenase (EC 1.8.1.3) to taurine. The rate limiting enzyme is CDO. In contrast to CGL, CDO activity is *up*regulated by increases in both cysteine and methionine concentration (Beetsch and Olson 1998; Ohta et al. 2000; Kwon and Stipanuk 2001; Tappaz 2004). Thus, the increase in methionine amount that was induced by inhibiting TNAP would increase CDO activity, leading to increased concentration of hypotaurine and taurine.

Again reversing perspectives, our results suggest that hypotaurine and taurine amounts decreased in SK-N-SH cells that proliferated and/or that differentiated into neurons without constraint (control condition. Interestingly, Sano et al. (2018) reported that differentiation of cultured PC12 cells is associated with a strong decline of intracellular taurine concentration (≥ 2 fold in comparison to age matched cultures).

These results are at variance with a number of studies dealing with taurine and brain development. Indeed, while our results suggest that intracellular taurine concentration decreases in cells that proliferate and differentiate, *in vivo* studies instead showed that taurine deficiency in kitten delays brain cell maturation and profoundly affects neuronal differentiation and neurons’ migration (Sturman et al. 1985; Palackal et al. 1986). Furthermore, *in vitro* studies showed that exogenously applied taurine may promote proliferation and migration of neuronal precursor cells as well as synaptogenesis (reviewed in Kilb and Fukuda 2017). Note that these effects were observed with rather high concentrations of extracellular taurine, typically 300 µM to 5 mM (e. g., Hernández-Benítez et al. 2010; Shivaraj et al. 2012; Kilb and Fukuda 2017).

It is quite unlikely that extracellular taurine had any influence in our experimental conditions. Our culture media included FBS. Taurine concentration in fetal calf serum ranges between ≈60 and ≈180 µM (Baetz et al. 1975). Yet, as FBS was used at 2% in the differentiation medium, the basal extracellular concentration of taurine would only be 1.2-3.6 µM. As there was very little basal taurine in our culture media, intracellular taurine would have to be excreted in high amounts to achieve an effective extracellular concentration. Yet release of taurine hardly occurs spontaneously (Kürzinger and Hamprecht 1981; Beetsch and Olson 1998) and extracellular taurine level *in vivo* is maintained at low level (≤25 µM; e.g.: Benveniste et al. 1984; Hagberg et al. 1987; Goda et al. 1998) by an efficient taurine transporter (Kürzinger and Hamprecht 1981; Tappaz 2004). Significant increases in extracellular brain taurine levels occurs only under rather deleterious conditions such as ischemia (e.g.: Benveniste et al. 1984; Hagberg et al. 1987; Goda et al. 1998) and osmotic challenges (Tappaz 2004; Ramírez-Guerrero et al. 2022). Furthermore, in the developmental studies mentioned above, extracellular taurine has been shown to exert its action by activating GABA_A_ and/or glycine receptors (Kilb and Fukuda 2017; Ramírez-Guerrero et al. 2022). However, the absence of GABA in the metabolome of SK-N-SH cells (current study) and the absence of functional GABA_A_ receptors in SH-SY5Y cells (Nikonorov et al. 2003) suggest that GABAergic signaling is not part of the signaling mechanisms used by these neuroblastoma cell lines; moreover, we have been unable to find a single study reporting on electrophysiological response through glycine receptor activation in either SK-N-SH or SH-SY5Y cells.

Thus, even if taurine was released in large enough amounts in our experimental conditions, it would not find GABA_A_ receptors and probably no glycine receptors to exert its neurogenesis promoting actions. We therefore come to the disappointing hypothesis that intracellular taurine accumulation in the presence of the TNAP inhibitor may have no functional role in preventing or promoting proliferation and differentiation of SK-N-SH D cells and may simply reflect the need for scavenging the excess in cysteine that may result from the increase in methionine level.

In summary, our study shows that TNAP is involved in the proliferation of SK-N-SH cells and in their differentiation into neurons. These results were obtained despite the presence of pyridoxine in the culture media, suggesting that TNAP fulfilled these roles in addition to its role in PLP metabolism. The proliferation/differentiation-promoting action of TNAP was accompanied by significant variations in the concentration of several metabolites, including metabolites related to lipid metabolism, proteinogenic amino acids and organosulfur compounds. The decrease in the amount of methionine in cells that proliferate and differentiate normally could play a determining role in the production of glutathione at the expense of taurine. The characterization of the signaling pathways involved in the neu-rogenesis-promoting action of TNAP deserves further investigation.

## Acknowledgments/Disclosures

Acknowledgments

The authors thank Christophe Prehaud for the gift of the SK-N-SH D cell line. They also thank Christophe Prehaud and Myriam Ermonval for their advice in setting up the SK-N-SH D cell culture.

## Authors’ contributions

### Conceptualization

AB, LGN, SB, VG, DM and CF. *Investigation*: AB and SB. *Formal analysis*: AB, LGN, DM, SB and VG. *Visualization*: LGN. *Writing – original draft preparation*: LGN. *Writing – review and editing*: AB, LGN, SB, VG, DM and CF.

### Funding sources

The research costs of this study were covered by CNRS, Claude Bernard-Lyon 1 University and University of Toulouse.

## Data Availability Statement

The data that support the findings of this study are available from the corresponding author upon reasonable request.

## Abbreviations

AP: alkaline phosphatase
CDO: cysteine dioxygenase
FBS: fetal bovine serum
GCL: glutamate-cysteine ligase
LDH: lactate dehydrogenase
MTT: 3-(4,5-dimethyl thiazol-2-yl)-2,5-diphenyltetrazolium bromide
PLP: pyridoxal phosphate
pNPP: p-nitrophenyl phosphate
RRID: Research resource identifier
SVZ: subventricular zone
TNAP: Tissue-nonspecific alkaline phosphatase
TSP: sodium 2,2,3,3-tetradeutero-3-trimethylsilylpropionate
VZ: ventricular zone

