## Supplemental material for "Tissue-nonspecific alkaline phosphatase promotes neuronal cell proliferation and differentiation: metabolomic reveals glutathione and taurine as molecular correlates"

### Supplementary figure: Figure S1

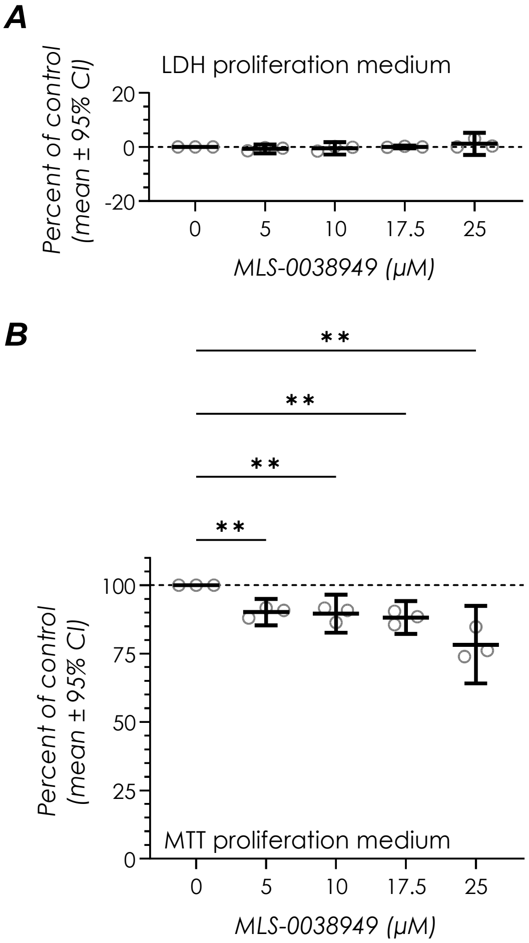


**Figure S1. Effects of inhibiting TNAP with MLS-0038949 on metabolic activity and cell viability in proliferation medium.** SK-N-SH D cells were seeded then grown for two days in the proliferation medium in the presence or absence of the TNAP inhibitor MLS-0038949. MLS-0038949 was applied at 5. 10. 17.5 and 25 µM. Dots in the graphs correspond to single experiments. Tests were made in duplicates in three experiments. The central horizontal line corresponds to the means and the upper and lower bars encompass the 95% confidence interval of the mean. **A.** LDH activity values were normalized such that a value of 100% corresponds to the toxicity induced by the complete cell lysis produced by Triton X100. and 0% to the LDH activity measured in control condition. We found no significant effect of MLS-0038949. whatever the dose. on LDH release (two-way ANOVA. $P = 0.14$). This implies that MLS-0038949 had no cytotoxic effect. whatever the dose. **B.** Cell metabolism was evaluated using the MTT test. using the same cultures as for the LDH test. The optical density of formazan generated from MTT was normalized to that measured in control condition (100%). MLS-0038949 induced a decrease in formazan level (two-way ANOVA. $P = 0.007$). Posthoc tests (Holm-Šídák's tests) indicated an overall decrease. in comparison to control. for all MLS-0038949 concentrations. Despite trends. we could not disclose a clear dose dependency in the action of MLS-0038949. At 25 µM. the TNAP inhibitor decreased the activity to 78% of control. a value very close to that obtained after two days in culture in differentiation medium (main text. Figure 2C). In both A and B. Geisser-Greenhouse correction was used to correct for lack of sphericity for the ANOVAs. Significance levels of posthoc tests are given by the number of asterisks: **: $P < 0.01$.

**Anne Briolay, Lionel G Nowak, Stéphane Balayssac, Veronique Gilard, David Magne, Caroline Fonta.**
*Tissue-nonspecific alkaline phosphatase promotes neuronal cell proliferation and differentiation: metabolomic discloses glutathione and taurine as molecular correlates*

#### Supplementary tables, part 1: full statistical report

**Table 1**
**Alkaline phosphatase activity (figure 1A): results of two-way ANOVA (mixed effect analysis)**

***ANOVA***

| | *Degrees of freedom* | *F value* | *P value* |
|---|---|---|---|
| Days in culture | 2. 18 | 3.526 | 0.0510 |
| Medium | 1. 9 | 14.21 | 0.0044 |
| Days in culture x medium | 2. 16 | 1.709 | 0.2125 |

***Holm-Šídák's multiple comparisons test***

| | *Adjusted P Value* |
|---|---|
| Day 1. Dif* vs. Prol** | 0.4327 |
| Day 2. Dif vs. Prol | 0.0024 |
| Day 3. Dif vs. Prol | 0.0270 |
| Prol. day 2 vs. day 1 | 0.9358 |
| Prol. day 3 vs. day 1 | 0.2474 |
| Prol. day 3 vs. day 2 | 0.2474 |
| Dif. day 2 vs. day 1 | 0.1316 |
| Dif. day 3 vs. day 1 | 0.0206 |
| Dif. day 3 vs. day 2 | 0.2979 |

*: Dif = differentiation medium; **: Prol = proliferation medium

**Table 2**
**Number of cells per well (figure 2A): results of two-way ANOVA**

***ANOVA***

| | *Degrees of freedom* | *F value* | *P value* |
|---|---|---|---|
| Days in culture | 1. 6 | 118.7 | <0.0001 |
| Treatment | 1. 6 | 14.21 | 0.0155 |
| Days in culture x treatment | 1. 6 | 19.77 | 0.0043 |

***Holm-Šídák's multiple comparisons test***

| | *Adjusted P Value* |
|---|---|
| Day 1. control vs. MLS | 0.7624 |
| Day 2. control vs. MLS | 0.0004 |
| Control. day 1 vs. day 2 | <0.0001 |
| MLS. day 1 vs. day 2 | 0.0038 |

**Table 3**
**Protein per well (figure 2B): results of two-way ANOVA**

**ANOVA**

| | *Degrees of freedom* | *F value* | *P value* |
|---|---|---|---|
| Days in culture | 1. 6 | 709.6 | <0.0001 |
| Treatment | 1. 6 | 10.48 | 0.0178 |
| Days in culture x treatment | 1. 6 | 40.17 | 0.0007 |

**Holm-Šídák's multiple comparisons test**

| | *Adjusted P Value* |
|---|---|
| Day 1. control vs. MLS | 0.3566 |
| Day 2. control vs. MLS | 0.0005 |
| Control. day 1 vs. day 2 | <0.0001 |
| MLS. day 1 vs. day 2 | <0.0001 |

**Table 4**
**MTT test (figure 2C): results of two-way ANOVA**

**ANOVA**

| | *Degrees of freedom* | *F value* | *P value* |
|---|---|---|---|
| Days in culture | 1. 3 | 94.91 | 0.0023 |
| Treatment | 2. 6 | 16.76 | 0.0035 |
| Days in culture x treatment | 2. 6 | 6.63 | 0.0302 |

**Holm-Šídák's multiple comparisons test**

| | *Adjusted P Value* |
|---|---|
| Day 1. control vs. MLS 10 µM | 0.5572 |
| Day 1. control vs. MLS 25 µM | 0.2091 |
| Day 1. MLS 10 µM vs. MLS 25 µM | 0.3235 |
| Day 2. control vs. MLS 10 µM | 0.0503 |
| Day 2. control vs. MLS 25 µM | 0.0021 |
| Day 2. MLS 10 µM vs. MLS 25 µM | 0.0503 |
| Control. day 1 vs. day 2 | 0.0001 |
| MLS 10 µM. day 1 vs. day 2 | 0.0004 |
| MLS 25 µM. day 1 vs. day 2 | 0.0011 |

**Table 5**
**LDH test (figure 2D): results of two-way ANOVA** (with Geisser-Greenhouse's correction)

**ANOVA**

| | *Degrees of freedom* | *F value* | *P value* |
|---|---|---|---|
| Days in culture | 1. 3 | 0.1442 | 0.7294 |
| Treatment | 1.610. 4.831 | 6.345 | 0.0477 |
| Days in culture x treatment | 1.303. 3.908 | 2.065 | 0.2343 |

**Holm-Šídák's multiple comparisons test**

| | *Adjusted P Value* |
|---|---|
| Day 1. control vs. MLS 10 µM | 0.3604 |
| Day 1. control vs. MLS 25 µM | 0.5302 |
| Day 1. MLS 10 µM vs. MLS 25 µM | 0.3129 |
| Day 2. control vs. MLS 10 µM | 0.9996 |
| Day 2. control vs. MLS 25 µM | 0.8537 |
| Day 2. MLS 10 µM vs. MLS 25 µM | 0.1845 |
| Control. day 1 vs. day 2 | (NA. mean dif. = 0) |
| MLS 10 µM. day 1 vs. day 2 | 0.2712 |
| MLS 25 µM. day 1 vs. day 2 | 0.9431 |

**Table 6**
**Number of neurites per cell (figure 3D): results of two-way ANOVA**

**ANOVA**

| | *Degrees of freedom* | *F value* | *P value* |
|---|---|---|---|
| Days in culture | 1. 156 | 5.159 | 0.0245 |
| Treatment | 1. 156 | 45.89 | <0.0001 |
| Days in culture x treatment | 1. 156 | 0.1887 | 0.6646 |

**Holm-Šídák's multiple comparisons test**

| | *Adjusted P Value* |
|---|---|
| Day 1. control vs. MLS | <0.0001 |
| Day 2. control vs. MLS | <0.0001 |
| Control. day 1 vs. day 2 | 0.1118 |
| MLS. day 1 vs. day 2 | 0.1959 |

**Table 7**
**Neurite lengths (figure 3E): results of nested t-test on log transform of the data**

| | *Degrees of freedom* | *t value* | *P value* |
|---|---|---|---|
| Day 1. control vs. MLS | 78 | 1.0930 | 0.2776 |
| Day 2. control vs. MLS | 78 | 0.1439 | 0.8859 |

**Table 8**
**Metabolomics data (figure 4): results of the two-tailed paired t-test**

| *Metabolite* | *Degrees of freedom* | *t value* | *P value* | *A posteriori power* |
|---|---|---|---|---|
| Acetate | 4 | 4.653 | 0.0096 | 0.93 |
| Alanine | 4 | 2.735 | 0.0522 | (0.55) |
| AMP | 4 | 0.02448 | 0.9816 | |
| Aspartate | 4 | 0.8747 | 0.4311 | |
| Choline | 4 | 2.066 | 0.1077 | |
| CMP | 4 | 0.5837 | 0.5907 | |
| Creatine | 4 | 0.5609 | 0.6048 | |
| Fumarate | 4 | 3.056 | 0.0378 | 0.63 |
| Glutamate | 4 | 2.589 | 0.0607 | (0.5) |
| Glutamine | 4 | 2.652 | 0.0569 | (0.52) |
| Glutathione | 4 | 3.457 | 0.0259 | 0.74 |
| Glycerophosphocholine | 4 | 3.638 | 0.0220 | 0.78 |
| Glycine | 4 | 3.662 | 0.0215 | 0.78 |
| GMP | 4 | 0.01987 | 0.9851 | |
| Histidine | 4 | 2.609 | 0.0595 | (0.52) |
| Hypotaurine | 4 | 6.172 | 0.0035 | 0.99 |
| Isoleucine | 4 | 3.443 | 0.0262 | 0.74 |
| Lactate | 4 | 0.3282 | 0.7592 | |
| Leucine | 4 | 3.456 | 0.0259 | 0.73 |
| Methionine | 4 | 3.714 | 0.0206 | 0.79 |
| NAD+ | 4 | 1.524 | 0.2023 | |
| Nicotinamide | 4 | 0.6323 | 0.5615 | |
| Phenylalanine | 4 | 1.166 | 0.3083 | |
| Phosphocholine | 4 | 1.611 | 0.1826 | |
| Phosphocreatine | 4 | 0.3881 | 0.7177 | |
| Proline | 4 | 3.913 | 0.0173 | 0.83 |
| Succinate | 4 | 0.2284 | 0.8306 | |
| Taurine | 4 | 4.418 | 0.0115 | 0.9 |
| Tyrosine | 4 | 0.5067 | 0.6390 | |
| UDP | 4 | 0.5771 | 0.5948 | |
| UMP | 4 | 3.747 | 0.0200 | 0.8 |
| Uridine | 4 | 2.31 | 0.0821 | |
| UTP | 4 | 2.754 | 0.0512 | (0.56) |
| Valine | 4 | 3.9 | 0.0175 | 0.83 |

The probability of making a type 1 error being set at 5%, either one or two of the significant differences reported here would be a false positive ($0.05 \times 34 = 1.7$). Yet significant differences were observed in a much larger number of comparisons (13/34 comparisons). The *a posteriori* power of the tests is presented for the *P* values that were < 0.05 and, between parenthesis, for the *P* values between between 0.05 and 0.06.
